# The anti-lipidemic drug simvastatin modulates Epigenetic Biomarkers in the Amphipod *Gammarus locusta*

**DOI:** 10.1101/2020.04.23.058248

**Authors:** Nélson Alves, Teresa Neuparth, Susana Barros, Miguel M. Santos

## Abstract

The adverse effects of certain environmental chemicals have been recently associated with epigenome’s modulation. Although the changes in the epigenetic signature are still not integrated into hazard and risk assessment, they are interesting candidates for linking environmental exposures to altered phenotypes given that these changes may be passed across multiple non-exposed generations. Here, we addressed the effects of simvastatin (SIM), one of the most prescribed human pharmaceuticals, in epigenetic regulators of the amphipod *Gammarus locusta*, as a proxy to support its integration in hazard and environmental risk assessment. SIM is a known modulator of epigenome in mammalian cell lines, and has been reported to impact *G. locusta* ecological endpoints at environmentally relevant levels. *G. locusta* juveniles were exposed to three SIM concentrations (0.32, 1.6 and 8 µg.L^-1^), for 15 days. The basal expression of selected epigenetic regulators was determined, along with the quantification of DNA methylation levels and the assessment of key ecological endpoints. Exposure to 0.32 and 8 µg.L^-1^ SIM induced significant downregulation of DNA methyltransferase1 (*dnmt1)*, concomitantly with Global DNA hypomethylation and impact on growth. Overall, this work is the first to validate the basal expression of key epigenetic regulators in a keystone marine crustacean, supporting the integration of epigenetic biomarkers into hazard assessment frameworks.

## 1. Introduction

Alterations in environmental conditions can trigger selection pressures in natural populations that experience conditions outside their physiological tolerances [1–3] and, therefore, influence the ability of species to survive and evolve [4]. Epigenetic variation is environmentally sensitive and can explain part of the organism’s responses, working as an adaptive response to the environmental changes [5,6,7,9]. However, the disruption of epigenetic machinery may lead to harmful effects in the organisms, which are potentially transmitted to the next generations [10–12]. Epigenetic inheritance has emerged as a rapidly growing field in the environmental sciences. It is well established that epigenetic mechanisms are highly responsive to external stimuli [6] and a wide range of environmental chemicals has been shown to generate specific epigenetic patterns in several organisms [13–16], highlighting its underlying role in the regulation of gene transcription [17,18].The modulation of these processes and its perpetuation can significantly impact the genetic and structural shape of population [19]. Therefore, epigenetic signatures are interesting candidates to link environmental exposures to altered phenotypes, providing new insights into the heritability of multigenerational exposure history [12,20–22]. Despite the fact that biomarkers of epigenetic modifications are still not integrated into hazard and risk assessment frameworks [12,23], a number of recent studies has suggested that risk assessment can benefit from the integration of chemical-induced epigenetic effects into toxicity testing strategies and hazard assessment [24–27].

Statins, such as simvastatin (SIM), are among the most prescribed human pharmaceuticals, known to reach the aquatic environments in increasing concentrations [28–31]. It is well established that, in vertebrates and arthropods, SIM disrupts the mevalonate pathway (MP) by inhibiting the enzyme 3-hydroxy-3-methylglutaryl-coenzyme A reductase (HMGR) [32,33]. However, whereas vertebrates obtain cholesterol from MP, cholesterol is not synthesized *de novo* in crustaceans; yet, they synthesize methylfarnesoate (MF) in the MP that has a central role in crustacean reproduction [34]. We have recently evaluated the chronic effects of SIM in the amphipod *Gammarus locusta* and our findings showed a significant impact on reproduction and growth at environmentally relevant concentrations [35]. Several recent studies have also reported growth, reproductive and embryonic development alterations in different metazoans under SIM exposure [35–41]. Although the underlying mechanism(s) of the observed SIM effects are poorly understood, previous studies with mammalian cell lines suggested that this pharmaceutical is able to modulate the regulation of the epigenome (e.g. DNA Methylation, histone acetylation and ncRNAs)[42–46]. Alterations in DNA methylation have previously been associated with downregulation or inhibition of DNA methyltransferase 1 (DNMT1), a critical protein responsible for maintenance of DNA methylation patterns [47,48]. Methylation of CpG islands, dense regions of CpG sites often located in gene promoters [49,50], is associated with gene repression [51,52]. The maintenance of methylation patterns is achieved by DNMT1 [53]. Despite of its major role in the maintenance of DNA methylation status, several evidences that DNMT1 cannot maintain the global DNA methylation by itself has been raised [54,55]. Several studies evaluated the effects of DNMT1 transcriptional changes and their downstream consequences. However, the upstream regulation of this critical enzyme is not commonly addressed. Epigenetic changes, definitive or transient, can influence and compromise the biological processes in the organisms [56]. Therefore, the integration of these proteins as potential biomarkers of epigenetic regulation is critical for an improvement of hazard and risk assessment. These biomarkers can potentially predict adverse outcome effects along the organism’s lifecycle [57,58], representing inherent, stable and robust mechanisms of memory of environmental exposures [59] and thus potentially providing an easy, sensitive and accurate tool to address changes after environmental exposures [60].

Following the findings presented above, in the present study we selected the human pharmaceutical SIM and the keystone amphipod *G. locusta*, to identify and validate the use of potential epigenetic biomarkers in ecotoxicological studies. The keystone marine species, *Gammarus locusta*, is an interesting test species due to its sensitivity to a wide variety of contaminants, easy culturing and its presence on the base of several trophic chains, highlighting its ecological relevance in aquatic ecosystems [35,61–66]. Furthermore, its short lifecycle is a great advantage for applying this species in epigenetic studies.

As a case study, we exposed *G. locusta* to environmentally relevant levels of SIM and a battery of epigenetic biomarkers (DNA methylation levels and the expression of key epigenetic regulators: DNA methyltransferase 1 - *dnmt1*, DNA methyltransferase 1-associated protein 1 - *dmap1*, E3-ubiquitin-protein ligase UHRF1 - *uhrf1*, Histone-Acetyltransferase 5 - *kat5* and Ubiquitin-Specific Peptidase 7 - *usp7*) were evaluated concomitantly with endpoints at individual-level (survival and growth).

## 2. Results

### 2.1. Basal Expression levels

This study is the first to characterize basal expression of epigenetic-related genes in the genus *Gammarus*. Figure 1A compares the gene transcription between several selected epigenetic regulators. Moreover, transcription level of the selected candidate housekeeping genes was also assessed (Figure 1B). All data were normalized to *clathrin. actin* was the most transcribed gene, expressing 239.9-fold more than the other housekeeping genes and *eif2* only being expressed 7-fold more, when both were compared with *clathrin*. *rpl13* and *gapdh* were transcribed 118.6 and 69.3 times more than c*lathrin*, respectively.

**Figure 1.**
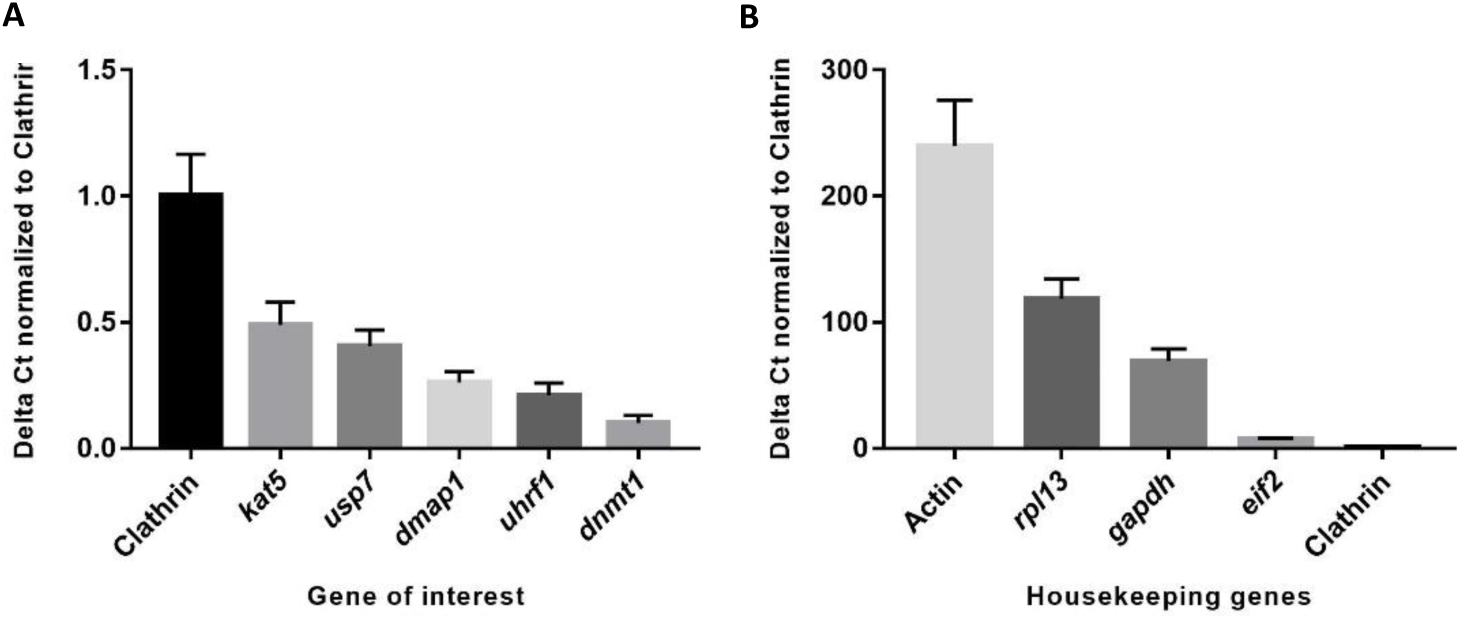
A) Basal expression of tested epigenetic regulators and B) validated housekeeping genes in *Gammarus locusta.* All data is presented in relative expression to *clathrin* (mean ± standard error). N=7-8.

In our assay, *dnmt1* (CT mean = 27.73) was the less expressed gene with 10-fold less transcripts in comparison to *clathrin*. *kat5* (CT mean = 24.89) and *usp7* (CT mean = 25.19) displayed similar transcription levels with 2.0 and 2.5-fold less transcripts than *clathrin*, respectively. *uhrf1* (CT mean = 26.27) and *dmap1* (CT mean = 25.75) were transcribed 4.7 and 3.8 times less than *clathrin* (Figure 1A).

### 2.2. Simvastatin as a case study

#### 2.2.1. Ecological Parameters

Figure 2A and B displays the mortality rate and the metasomatic length of *G. locusta* after 15-days of SIM exposure. The exposure of *G. locusta* to the three SIM concentrations (0.32 to 8 µgL^-1^) did not induce any statistically significant differences in mortality. On the other hand, a dose-dependent increase in metasomatic growth was observed with statistically significant differences at 1.6 and 8 µg.L^-1^. It should be noted that at the end of exposure amphipods were immature without any displayed secondary sexual characters.

**Figure 2.**
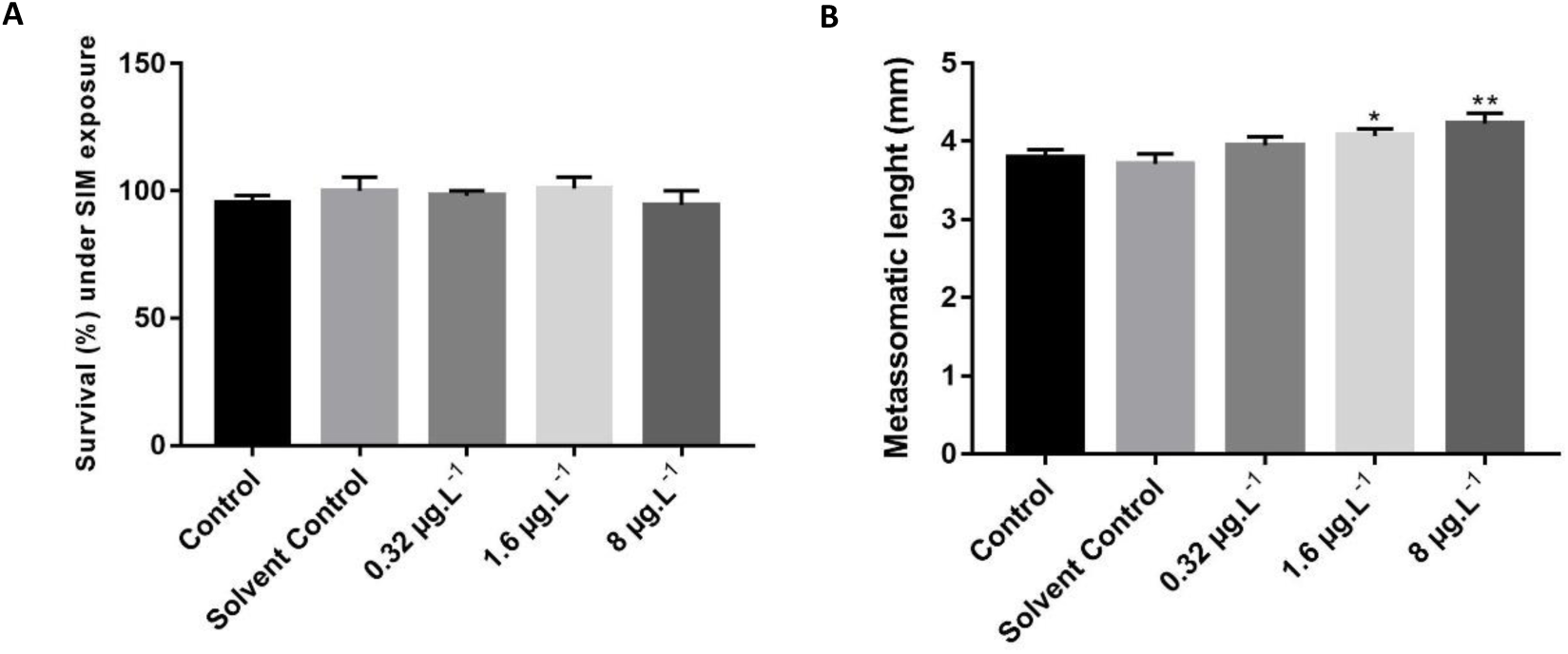
*A)* Survival of *Gammarus locusta* after 15-days of exposure to several SIM concentrations. Data were normalized to control survival and presented as mean ± standard error. N=2 B) Metasomatic length of *Gammarus locusta* after 15-days of exposure to several SIM concentrations. Data were presented as mean ± standard error. N= 34-35. Significant statistical differences, p<0.05 and p<0.01, are highlighted with asterisks (*)(**), respectively.

#### 3.2.2. Gene transcription of epigenetic regulators

An upregulation was observed in the gene transcription of the housekeeping genes at 8 µg.L^-1^ of SIM. Since all the housekeeping genes tested (*actin-C*, *eif2, rpl13, gapdh, clathrin*) presented the same pattern for all SIM tested concentrations (Figure 3), it is plausible to hypothesize that *G. locusta*, at the highest SIM concentration (8 µg.L^-1^), was experiencing an increase in the overall metabolism. A recent study calculated the stability coefficient of several candidate reference genes in *Gammarus fossarum* using five different algorithms and showed that *clathrin* and *gapdh* were the two most stable genes in this species [67]. Given that the reference genes must show a stable transcriptional pattern across treatments [68], the qRT-PCR data was normalized here with *clathrin* and *gaphd*, the genes with the most similar transcription levels to the genes of interest. Yet, all tested reference genes were stable across treatments, with the exception of the 8 µg.L^-1^ group.

**Figure 3.**
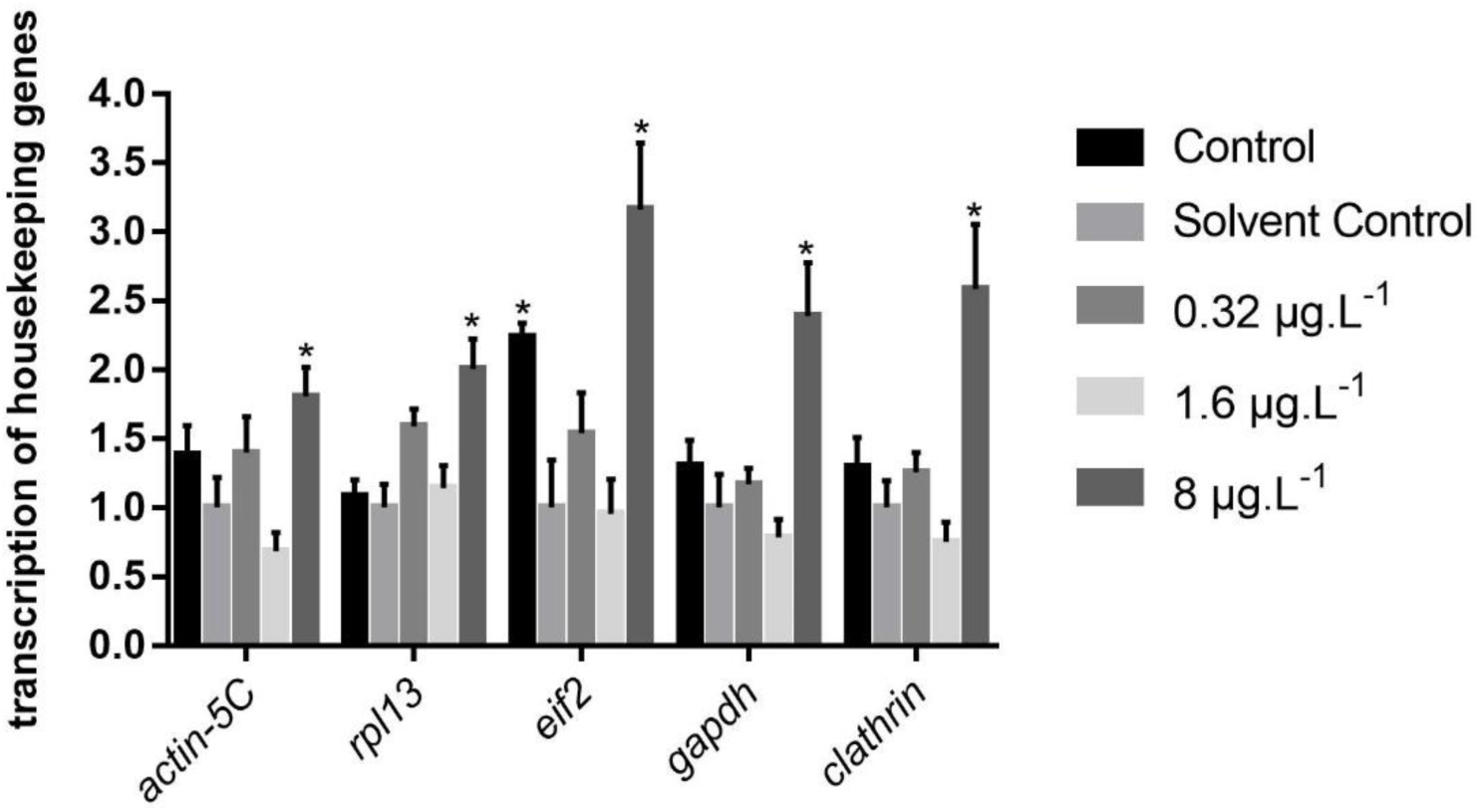
Pattern of mRNA transcription levels of candidate reference genes after an exposure for 15-days to SIM. Data (Mean ± Standard Error) are normalized to transcription levels of Solvent Control. Significant statistical differences (p<0.05) are highlighted with asterisks (*). N =7-8.

Concerning the genes of interest, SIM induced significant changes in the transcription pattern of *dnmt1* and *dmap1.* For *dnmt1*, a significant decrease in transcription was observed at 0.32 µg.L^1^ and 8 µg.L^-1^. In contrast, a significant upregulation of *dmap1* transcripts was observed at 1.6 µg.L^-1^ SIM (Figure 4).

**Figure 4.**
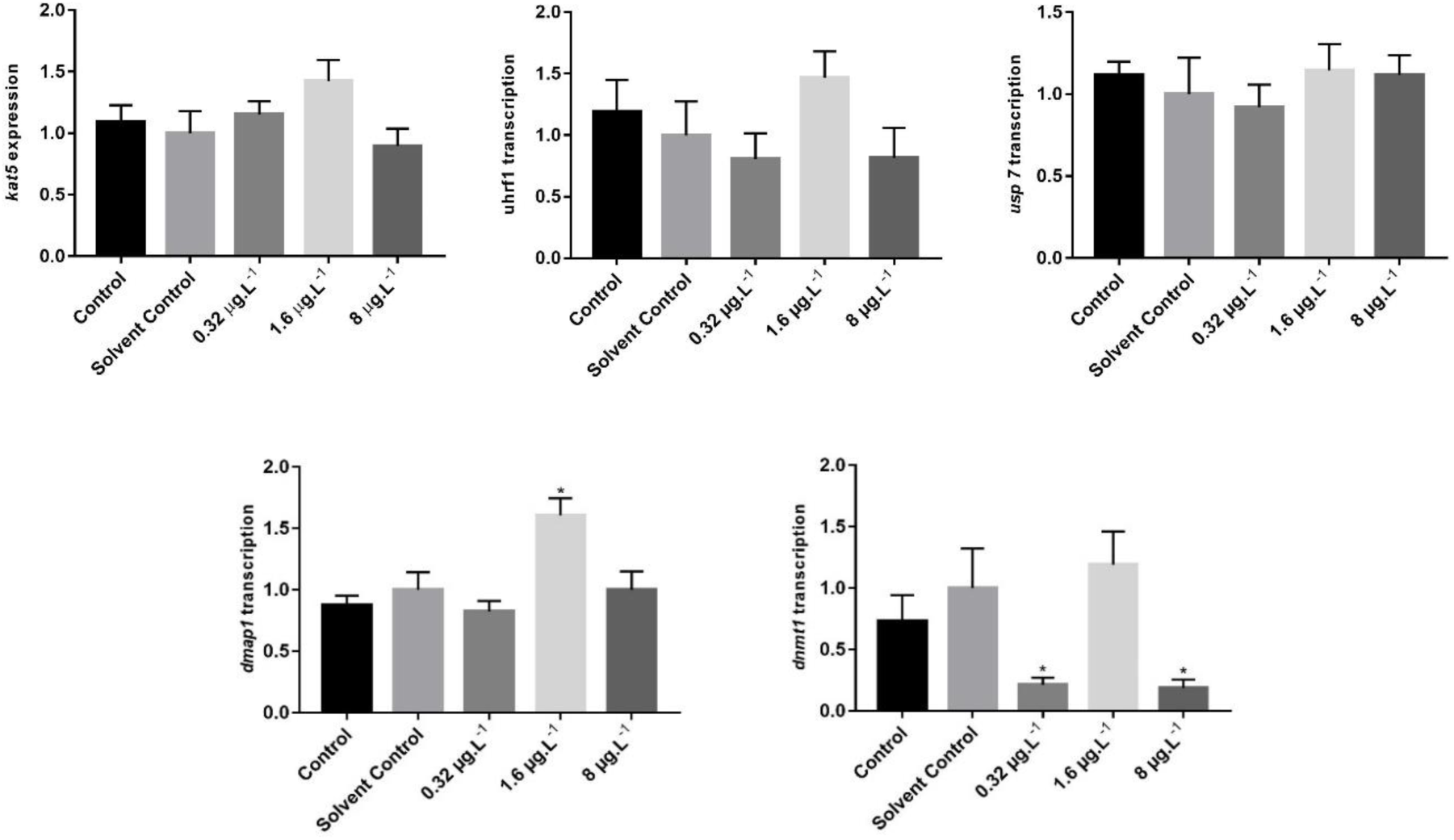
Pattern of mRNA transcription levels of studied epigenetic regulators after an exposure for 15-days to SIM. Data (Mean ± Standard Error) are normalized to *gapdh* and *clathrin* transcription levels and presented as fold-changes relative to the Solvent Control. Significant statistical differences (p<0.05) are highlighted with asterisks (*). N= 7-8.

**Figure 5.**
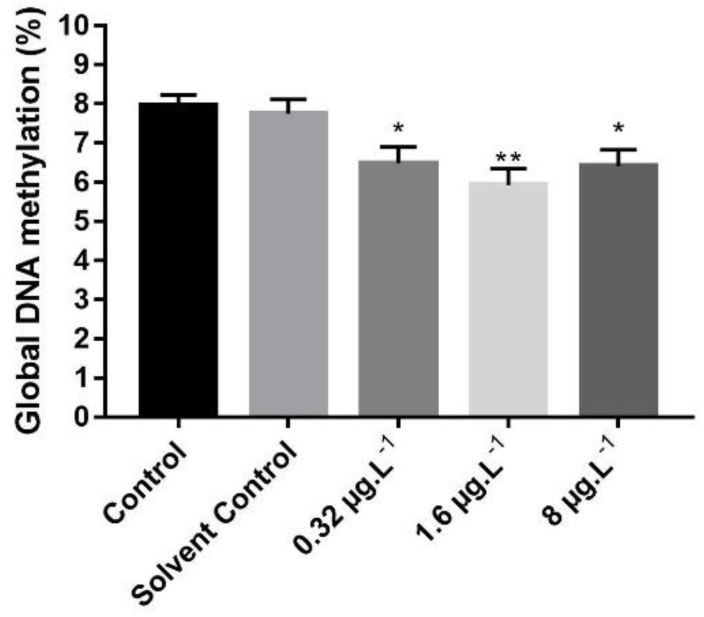
Percentage of Global DNA Methylation (Mean ± Standard Error) after 15 days of SIM exposure. Significant statistical differences are highlighted with asterisks (*) for p<0.05 and (**) for p<0.01. N= 8.

#### 2.2.3. Global DNA methylation

A significant decrease in global DNA methylation levels for the three tested treatments was observed. These data are in accordance with the data obtained for *dnmt1* transcripts at 0.32 µg.L^-1^ and 8 µg.L^-1^.

## 3. Discussion

In the last decade, a growing scientific interest on transgenerational epigenetic inheritance has emerged [69–71]. However, despite the increasing number of studies showing transgenerational epigenetic impacts, the integration of epigenetic markers in the frame of hazard and risk assessment is still not a reality. Currently, environmental chemical hazard and risk assessment frameworks rely mostly on hazard identification, dose−response studies, exposure assessment and risk characterization, often using a tier approach in four main categories (tier 1-4), without addressing transgenerational and epigenetic effects [12,72].

Understanding the long-term effects of epigenetic changes can improve our knowledge about the adaptive regulation and/or disruptive responses, addressing the concerns about how these effects may modulate the populations’ evolution [73]. Therefore, the identification and validation of biomarkers of epigenetic modifications can be of great interest as early warning tools in risk assessment frameworks to mitigate the risks of environmental exposures [12,23,74,75].

DNA methylation is a process relatively stable. In mammals, erasure of this mark can be achieved by an active or passive demethylation. Active demethylation involves the presence of enzymes called Ten-Eleven Translocases (TETs) that progressively oxidize 5-mC and transforms it in other bases more unstable. On the other hand, passive demethylation occurs in DNA replication. If the process of DNA methylation is not effective, a passive demethylation occurs concomitantly with the synthesis of new DNA strands. Therefore, the accurate maintenance of DNA methylation is critical to faithfully propagate specific patterns along individuals and generations [76,77].

The absence of any genes-encoding TETs in the transcriptome of *G. locusta* (unpublished data) suggest the importance of passive demethylation process and the putative role of *dnmt1* in maintaining the patterns of cytosine methylation in this species. Here, we evaluated the gene transcription levels of several epigenetic regulators and the levels of global DNA methylation, in *G. locusta* after 15 days of exposure to SIM, as a proxy to validate potential biomarkers for early detection of adverse effects. In the current study we observed a significant downregulation of *dnmt1* in *G. locusta* exposed to SIM concentrations of 0.32 and 8 µg.L^-1^.These findings integrate well with similar observations reported in mammalians models under SIM exposure [47,78]. In cancer cell lines, Karlic *et al*. [78] reported the downregulation of DNMT1 mRNA transcription in response to SIM. Together, these results give further support to the use of *dnmt1* gene as an early warning biomarker of epigenome modifications.

In mammalian genomes, downregulation of DNMT1, the protein responsible for DNA maintenance of methylation process, can lead to global DNA hypomethylation, that is frequently associated with an increase of global gene transcription [79]. Several studies with SIM indicated upregulation of Krüppel-like factor 2 (KLF2) genes [80], sterol regulatory element-binding protein (SREBP) [81], aryl hydrocarbon receptor (AhR), pregnane X receptor (PXR) and downstream pathways as CYP1a, CYP3a [82,83] and peroxisome proliferator-activated receptor (PPARs) [84,85]. A downregulation of several proteins is also frequently associated with epigenetic machinery. For example, enhancer of zeste homolog-2 (EZH2), a histone lysine methylase [86] and DNMT1 [47] were reported as downregulated proteins in human colorectal cancer (CRC) cell line after SIM exposure. This appears to support the hypothesis that SIM is able to induce global DNA hypomethylation. To confirm this hypothesis, in the present study we quantified the *G. locusta* global DNA methylation levels after SIM exposure. Our data revealed a global DNA hypomethylation at all SIM treatments, thus supporting the downregulation of *dnmt1* at 0.32 µg.L^-1^ and 8 µg.L^-1^.

As the *dnmt1* expression was not affected by SIM exposure at 1.6 µg.L^-1^, the global DNA hypomethylation observed at this SIM concentration could also be associated with other genes. The upregulation of *dmap1* may mediate the observed global DNA hypomethylation response. Studies in human cell lines and mice revealed that DNA hypomethylation is associated with DNA instability [87,88]. Negishi *et al.* [89] demonstrated that DMAP1 is requited to the maintenance of DNA integrity by mediating the involvement of the essential repairing machinery to DNA regions with double-strand breaks, ensuring a faithful DNA repair and replication [90]. In order to prevent this instability, a switch between chromatin status occurs, in a fluid manner, creating feedback loops in chromatin condensation and allowing the adaptation to external stimuli [91]. This switch is accomplished by the modulation of acetylation of lysine 16 in histone H4 (H4K16ac), a DMAP1-induced, acting in inter-nucleosome interaction and as a scaffold for other transcription factors and chromatin-modifying proteins [92,93]. A growing body of evidences suggests that statins can promote the acetylation of H3 and H4 histones [45,78,94]. Thus, the increase of *dmap1* transcription may be directly related with the increasing transcription of several genes or through the negative feedback required for the protection of the genome. However, to clarify this mechanism, future studies should evaluate the interplay between these proteins and their binding partners in specific sites of chromatin and additional insights into histone modifications should be disclosed.

In this work, we also measured the *G. locusta* metasomatic length, which for control animals were similar to the basal length reported for the same age class [63]. This fact allowed the validation and the use of this endpoint. SIM exposure induced a dose-response curve with an increase of metasomatic length in *G. locusta*, at the two highest tested concentrations. Dahl *et al.* [95] reported that exposure of a copepod, *Nitocra spinipes*, from 24-hours Nauplii until the3^rd^ copepodite stage, induced a significant increase of body length at 1.6 µg.L^-1^ SIM. We hypothesize that the increase of metasomatic length after SIM exposure may reflect an adaptive phenomenon after 15-days of SIM exposure and could represent an hormesis response, known to trigger the overall increase in metabolism. Interestingly, Neuparth *et al*. [35] observed a decrease metasomatic length in *G. locusta* after 36-days exposure under 0.32 µg.L^-1^, 1.6 µg.L^-1^ and 8 µg.L^-1^ SIM conditions. If the SIM exposure is prolonged beyond the 15 days, a subsequent instability in critical biological processes may override the observed stress-induced responses and lead to the adverse effects reported in Neuparth *et al*. [35]. In the study of Neuparth *et al*. [35], *G. locusta* was exposed to SIM during 36-days and lethality was observed at 8 µg.L^-1^. Moreover, *G. locusta* growth was impaired in a sex-dependent manner with a higher sensitivity in females, whose reproductive performance was reduced or completely hampered.

The adverse effects of SIM exposure [35] and the disruption of critical epigenetic machinery, can lead to the perpetuation of instability and adversity, resulting in disruption of critical biological processes in *G. locusta.* These findings suggest that the epigenetic disruption should be included in further studies performed to evaluate the multigenerational and transgenerational effects of SIM.

Furthermore, compounds with the potential to inhibit MP have been shown to act in one carbon metabolism [96–98]. This pathway is involved in S-adenosyl methionine (SAM) production, the methyl donor for DNA methylation reactions. The inhibition of this pathway induces a decreasing in SAM levels and may mediate a potential demethylation [99]. These previous studies give further support to the findings presented here.

Concerning other tested genes in the present study, *uhrf1*, *kat5* and *usp7*, showed a stable pattern along the different concentrations of SIM. The absence of changes in the transcription levels of several genes related with epigenetic could lead to the hypothesis that these may not work as early warning markers [12]. Despite the transcription of several selected genes were not altered after SIM exposure, these genes (*uhrf1, kat5 and usp7)* have essential functions in epigenetic machinery; and therefore could be promising biomarkers in the context of other environmental pollutants. Future work should evaluate the transcription of these genes in the context of exposure to other chemicals reported to modulate the regulation of the epigenome.

Overall, this study propose new epigenetic biomarkers in *G. locusta* as a proxy to improve hazard and risk assessment. The disruption of epigenetic machinery is the main process responsible for the propagation of adversity over generations. The inheritance of methylation profiles across DNA replication rounds is propagated through the activity of DNMT1. The disruption of DNMT1 levels can modify the methylation of DNA regions, leading to the transfer of this adverse effect throughout mitotic and meiotic processes by the disruption of methylation patterns in germline. Thereby these effects can be potentially propagated to subsequent generations, including non-exposed. The inclusion of epigenetic biomarkers proposed in this study in future hazard and environmental risk assessment and in the environmental management framework guidelines may provide, along with other data, important insights into potential transgenerational inheritance after contaminants exposure. The findings of this work, together with previous studies, supports the integration of DNMT1, DMAP1 and the global DNA methylation as a biomarker in hazard and environmental risk assessment.

## 4. Material and Methods

### 4.1. Amphipods Culture System

A permanent laboratory culture system of *G. locusta* is established at Interdisciplinary Centre of Marine and Environmental Research – CIIMAR. The amphipods were maintained at 20°C and 33‰ salinity with photoperiod set to 16h:8h (light:darkness), following the methodology described in detail by Neuparth *et al.* [63]. Fine to medium sand with 1 cm layer and small stones were placed in the bottom of the aquaria to assure the optimal raising conditions for *G. locusta* and simulate natural habitat. Aeration was established with plastic tips, allowing constant bubbling. *Ulva sp.* and sediments were collected in Aguda beach (41°3’6"N 8°39’21"W), *Vila Nova de Gaia*, in a site devoid of direct contamination source. The culture system is partially renewed once per year with organisms collected in Sado Estuary (38° 31’14"N 8°53’32"W).

### 4.2. Experimental System

The experimental design followed the conditions described in Neuparth *et al.* [35]. Briefly, the bioassay was conducted, during 15 days, in 7 L aquaria housing 60 juvenile (immature) amphipods with 10-days old in a semi-static system with five exposure conditions, two controls (0.45µm-filtered natural seawater and a solvent control with 0.0005% acetone) and three simvastatin treatments (0.32 µg.L^-1^, 1.6 µg.L^-1^ and 8 µg.L^-1^) with 2 replicates. Simvastatin (CAS number 79902-63-9, ≥ 97%) was purchased from Sigma-Aldrich and was prepared in acetone. Acetone percentage in all aquaria except natural seawater control was 0.0005%. The concentrations of SIM selected for the present study were based upon our previous research [35], which observed a severe impact on *G. locusta* reproduction at the highest levels selected, and data from environmental levels in aquatic ecosystems. The experimental apparatus was checked daily to ensure feeding needs, aeration and removal of dead animals, if any. Total water renewals were performed every two days, together with new test solutions applied to the aquaria. Throughout the experiment, amphipods were fed *ad libitum* with *Ulva sp.* The experimental conditions were similar to the culture system described above. Due to the high stability of SIM in water revealed in our previous studies [35,41], that used similar experimental design to that here reported, the actual concentrations of SIM were not monitored in the present study. At the end of exposure, survival and metasomatic length was measured, according to the methods described in Neuparth *et al.* [63]. Finally, amphipods were individually preserved in RNAlater^®^ to ensure RNA/DNA quality and then stored at −80°C until further use.

### 4.3. RNA isolation and cDNA synthesis

Total RNA from eight immature individual amphipods was extracted with NZYol^®^ (Nzytech, Portugal) together with the illustra^TM^ RNAspin Mini RNA Isolation Kit (GE Healthcare, United Kingdom), following the manufacturer’s instructions. RNA was quantified with a Take3TM on a microplate reader (Biotech Synergy HT) coupled with the software Gen5 (version 2.0). RNA quality and quantity were verified by electrophoresis in 1.5% agarose gel and by measurement of the ratio of optical density at λ260/280 nm. Synthesis of cDNA was performed with NZY First-Strand cDNA Synthesis Kit (nzytech^®^, Portugal), using 1 μg of total RNA in a total volume of 20 µL per reaction, as manufacturer’s instructions. At the end of the conversion, cDNA was stored at −20°C until qRT-PCR analysis.

### 4.4. Primers design and validation

The sequences of *G. locusta* used in this work were extracted from our *G*. *locusta* transcriptome (data not shown). *Hyalella* a*zteca* was used as a reference species to find genes of interest. Primers were designed in Primer-Blast from NCBI, a Primer3plus tool coupled to NCBI Blast [100]. Primers specificity was tested in range 60 ± 2.0°C. Amplification with Phusion^®^ High-Fidelity PCR Master Mix (Thermofischer, Portugal), was performed according Manufacturer’s Instructions, in a 20 µL total volume reaction. For each tested gene, product identity was confirmed by cloning in pGEM^®^-T Easy Vector Systems (Promega Corporation, USA), followed by Sanger sequencing. The genes selected for the present study were the following: 1) *dnmt1*, the methyltransferase critical for the maintenance of DNA methylation [53], 2) *uhrf1*, a central hub for maintenance of DNA methylation and a pivotal regulator of crosstalk between DNA methylation and histone modifications [101,102], 3) *dmap1*, a transcriptional co-repressor associated with DNMT1 *and* histone deacetylase 2 (HDAC2) in replication foci during S phase [103], 4) *usp7*, a deubiquitinating enzyme that prevents proteasomal degradation of UHRF1/DNMT1 complex [104,105] and 5) *kat5*, a histone acetyltransferase that induces USP7/DNMT1 destabilization and proteasomal degradation [106,107].

### 4.5. Real Time Quantitative-Polymerase Chain Reaction (qRT-PCR)

qRT-PCR was used to evaluate the gene transcription of *dnmt1, dmap1, uhrf1, usp7 and kat5*. Validated primers and fluorescence-based quantitative real time PCR (qRT-PCR) conditions are described in Table 1. Reactions were performed as Manufacturer’s instructions in a total volume of 10 μL. 400nM of each primer and 10ng of cDNA were used per well. All samples were assessed in duplicate. PCR reactions efficiency was determined by standard curves performed with six serial dilutions of the cDNA pool of all samples (from 0.0064 to 20 ng of cDNA) [41,108]. Reactions were conducted in a Mastercycler^®^ep realplex (Eppendorf, Spain). PCR conditions were as follows: 95°C for two minutes, followed by 35 cycles of 95°C for five seconds and combined annealing and extension at 58-62°C, for 25 seconds. Then, a melting curve analysis was performed (from 55°C to 95°C) to confirm specificity of products obtained. The presence of contaminations and potential by-products of PCR such as primers dimers were excluded by concomitant running of no template controls. PCR products were analysed by electrophoresis in 2% agarose gel to check the presence of a single expected band. Product’s identity was confirmed by sanger sequencing. Relative change in transcription abundance of target genes was normalized to *gapdh* and *clathrin* and calculated using the Livak method [109]. Expression levels were normalized to solvent control mean and data were expressed as fold changes relative to this group.

**Table 1.**
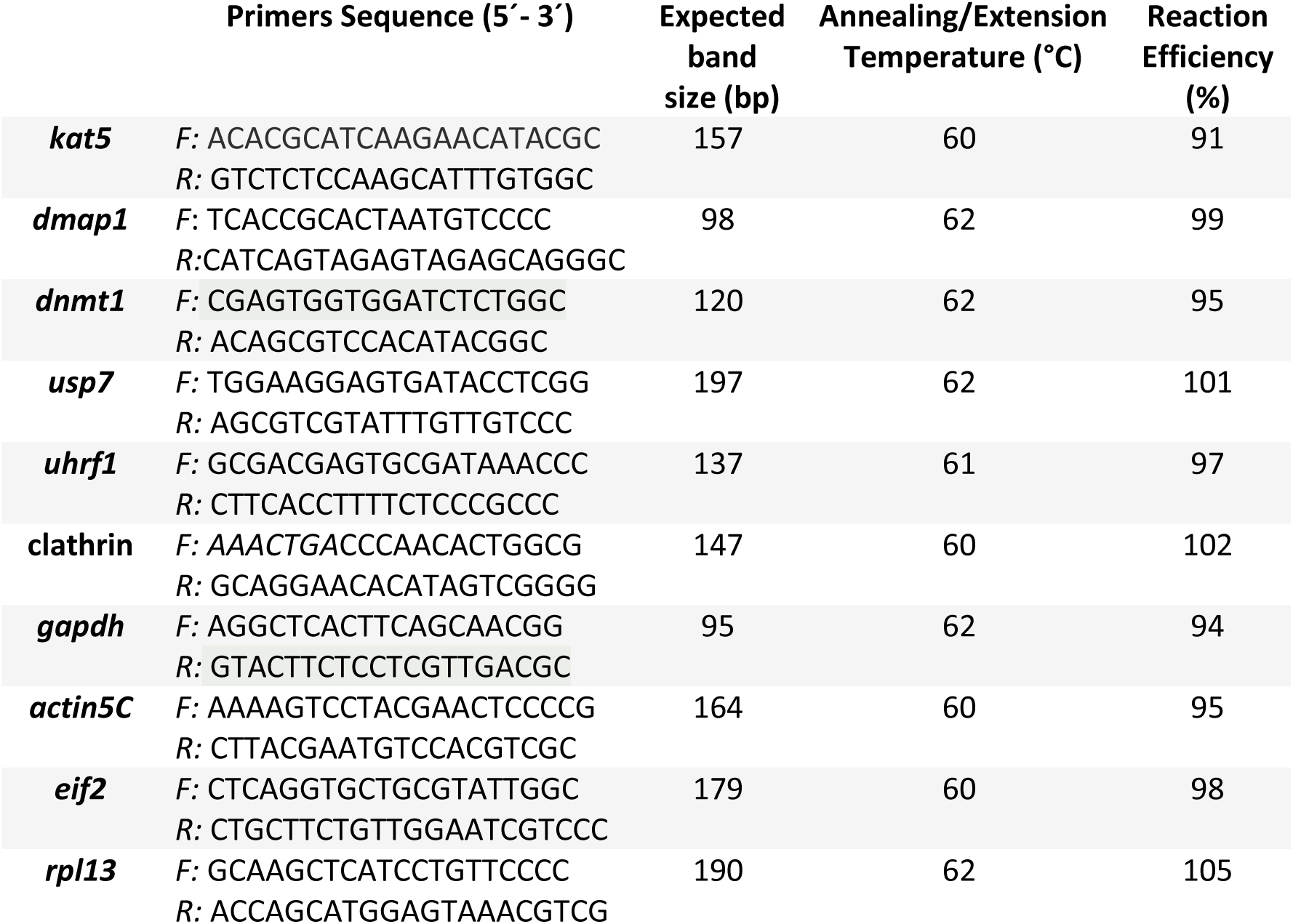
Sequences of primers (5’- 3’) used for evaluation of gene transcription as well as conditions used in amplification reaction. F - Forward Primer; R – Reverse Primer.

### 4.6. DNA isolation

Genomic DNA was isolated from eight individual amphipods with NZY Tissue gDNA Isolation Kit from nzytech® following the manufacturer’s instructions. A homogenization step was introduced in pre-lysis samples to facilitate the DNA extraction. DNA was quantified with a Take3^TM^ on a microplate reader (Biotech Synergy HT) coupled with the software Gen5 (version 2.0) and the DNA purity was assessed by measurement of the ratio of optical density at λ260/280 nm. Pure DNA was stored at −20°C until global DNA methylation quantification.

### 4.7. Quantification of global DNA methylation

Global DNA methylation was determined in eight DNA samples per treatment using the MethylFlash Global DNA Methylation (5-mC) ELISA Easy Kit (Colorimetric) from Epigentek® according to the manufacturer’s instructions. 100 ng of each DNA sample in duplicate was employed.

### 4.8. Statistical Analysis

Data obtained from this study were checked for homogeneity of variances (Levene’s test) and normality (Kolmogorov-Smirnov test). Data were transformed when one of these assumptions was not validated. Data were then analysed by one-way ANOVA. Post-hoc comparisons were carried out using Fisher’s least significant difference (LSD) test. Significant statistical differences were set as p<0.05. All treatments were compared with solvent control group. Statistica 13 (Statsoft, USA) was used to compute all analysis.

## Author Contributions

All experiments were designed by N.A., M.M.S. and T.N., and performed by N.A. and S.B. N.A. drafted the manuscript. S.B., T.N. and M.M.S. critically revised the manuscript. T.N. and M.M.S supervised all the work. All authors have read and agreed to the published version of the manuscript.

